# Evolutionary Response to Resource Deprivation: Parallelism and Nonmonotonicity

**DOI:** 10.1101/865584

**Authors:** Megan G. Behringer, Wei-Chin Ho, Samuel F. Miller, John C. Meraz, Gwyneth F. Boyer, Michael Lynch

**Author notes:** These authors contributed equally to this work (Megan G. Behringer, Wei-Chin Ho). Please direct correspondence to: Megan G. Behringer, Wei-Chin Ho.

## Abstract

Establishing reliable frameworks for predicting unknown outcomes from empirical observations is of great interest to ecologists and evolutionary biologists. Strong predictability in evolutionary responses has been previously demonstrated by the repeated observation of similar phenotypes or genotypes across multiple natural or experimental populations in analogous environments. However, the degree to which evolutionary outcomes can be predicted across environmental gradients, or in fluctuating environments, remains largely unexplored. Presumably, the phenotypic evolution in an intermediate environment could be interpolated from the evolved phenotypes observed in two extreme environments, but this assumption remains to be fully tested. Here, we report on the experimental evolution of *Escherichia coli* under three nutritional transfer periods: every day, every 10 days, and every 100 days, representing increasing severity in feast/famine cycles. After 900 days of experimental evolution, populations experiencing intermediate durations of starvation had evolved longer times to reach maximum growth rate, smaller colony sizes, higher biofilm formation, and higher mutation rates than populations evolving in the other environmental extremes. Because the intermediately starved populations exhibit significantly high molecular parallelism, these distinct phenotypes are likely due to non-monotonic deterministic forces instead of increased stochastic forces commonly associated with fluctuating environments. Our results demonstrate novel complexities associated with evolutionary predictability across environmental gradients and highlight the risk of using interpolation in evolutionary biology.

---

Evolution is driven by both deterministic (e.g., natural selection) and stochastic (e.g., genetic drift or environmental uncertainty) forces ^1–8^. Across analogous environments, similar phenotypes or genotypes are commonly observed suggesting strong deterministic forces and a potential for evolutionary outcomes to be highly predictable ^8–12^. Experimental evaluation suggests that higher environmental similarity leads to higher evolutionary predictability or parallelism, as two populations are more likely to have similar derived phenotypes and genotypes when evolved in more similar environments ^13–15^. Despite several empirical tests of evolutionary predictability across multiple distinct constant environments, there is still a lack of sufficient examination across fluctuating environments that differ both in the intensity of the stress imposed and the periodicity of fluctuations. Generating such an understanding of evolutionary predictability across fluctuating environments is of particular importance, as they are more representative of what organisms often face in nature.

One set of experimental conditions with significant biological relevance is feast/famine cycles. Many microbial populations, including facultative pathogens with clinical or agricultural importance, frequently encounter environments with fluctuating resource availability ^16,17^, *Escherichia coli*, for instance, has a broad habitat as an opportunistic pathogen, ranging from soil and wastewater to the lower gut, often oscillating between feast and famine ^18–20^. Prior studies have demonstrated that *E. coli* populations are capable of surviving periods of prolonged starvation, during which they accumulate mutations in the genes *rpoS* ^21^ and *lrp* ^22^ that confer a phenotype known as growth advantage in stationary phase (GASP)^23–25^. Other phenotypes observed in *E. coli* populations during resource limitation have included temporarily increased mutation rates ^26,27^ and the evolution of trade-offs between maximum growth rate and sustainable growth ^28^. Again, an understanding of how these genetic and phenotypic responses scale across different durations of resource limitation is still lacking. Thus, conclusions that the phenotypic response to repeated starvation, or any fluctuating selective pressure, scales progressively with an increasing duration of stress is premature.

To address this issue, we surveyed the evolutionary response of *E. coli* populations along a selective gradient of three resource replenishment cycles: every day, every 10 days, and every 100 days, and determined if constraints on the predictability of genotypes and phenotypes were associated with repeated long-term starvation. If, for example, evolutionary predictability is highly related to environmental similarity, phenotypic evolution in the intermediate environment (10-days) should be predictable from the evolved phenotypes observed in the two environmental extremes (1- and 100-days), i.e., the plot of phenotypic values against the environmental gradient should be monotonic (a continuously increasing or decreasing function). As different molecular and phenotypic traits have the potential to reveal distinct patterns in evolutionary responses, we focus on evolved changes in growth dynamics, mutation rate, genomic evolution rate, and evidence of subpopulation structure. We also examine the parallelism of derived mutations in populations evolving under the same or different nutrient limitation conditions.

## Results

To establish the evolution experiment, ancestral lines with two different initial genetic backgrounds were used: a wild-type strain (WT) and a WT-derived strain with impaired methyl-directed mismatch repair (MMR-) wherein an engineered deletion of *mutL* yields a ~150× increase in the single nucleotide mutation rate ^29^. Three distinct resource-limitation cycle conditions (1-day, 10-days, and 100-days between transfers into fresh media) were deployed, and each combination of genetic background and resource-limitation cycle condition was replicated in eight parallel experiments, resulting in 48 experimental populations (8×3×2). Thus, this design allows us to examine the parallelism within each treatment combination, as well as the divergence between treatments. After 900 days of experimental evolution, changes in growth behaviours were evaluated. Notably, populations subjected to 10-day cycles of resource limitation take significantly longer to reach their maximum growth rate (*T*_max_) than ancestral, 1- and 100-day populations (**Fig. 1a**). In addition, 10-day populations present significantly smaller colonies on LB agar than ancestral, 1-day, and 100-day populations (**Fig. 1b-e**).

**Figure 1.**
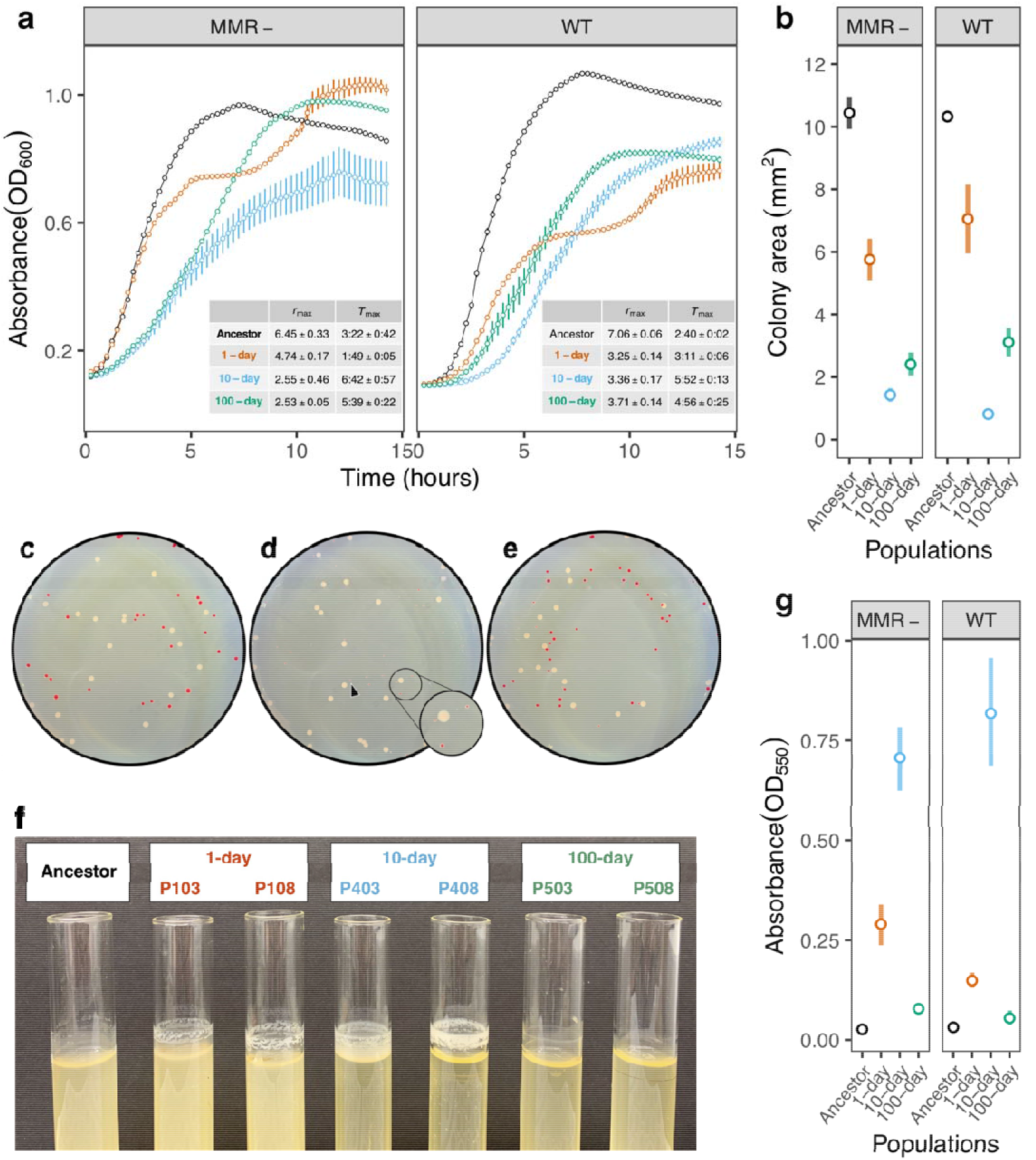
Phenotypic divergence of evolved lines after 900 days. Each resource-limitation cycle/genetic background combination is represented by two populations; 1-day (orange) - WT: P103, P108; MMR-: P106, P111; 10-day (blue) - WT: P403, P408; MMR-: P406, P411; 100-day (green) - WT: P503, P508; MMR-: P506, P511. Significant differences in growth phenotypes were determined by two-sided, pairwise Welch’s t-tests adjusted by FDR and all error bars represent ± s.e.m. **(a)** Growth curves of ancestor (black) and evolved populations cultured in LB broth at 37 °C. Circles denote average absorbance at 600 nm measured every 15 minutes for 14 hours. Nested table displays the averages ± s.e.m. for maximum growth rate (*r*_max_) and time to maximum growth rate (*T*_max_). *T*_max_ is presented in units of time (hours:minutes). Populations evolving to 10-day resource-limitation cycles take significantly longer to reach *T_max_* (WT: *t_anc_* = 14.91, *P_anc_* = 4.5 x 10^−10^; *t_1-day_* = 11.65, *P_1-day_* = 4.6 x 10^−10^; *t_100-day_* = 1.99, *P_100-day_* = 0.057; MMR-: *t_anc_*= 7.24, *P_anc_* = 2.5 x 10^−5^; *t_1-day_* = 7.81, *P_1-day_* = 8.4 x 10^−6^; *t_100-day_* = 1.87, *P_100-day_* = 0.073;). **(b)** Average colony sizes of evolved populations and ancestors after 24 h of growth on LB agar. Populations evolving to 10-day resource-limitation cycles produce significantly smaller colonies (WT: *t_anc_* = 44.99, *P_anc_* < 1 x10^−16^; *t_1-day_* = 5.56, *P_1-day_* = 3.3 x 10^−4^; *t_100-day_* = 4.85, *P_100-day_* = 5.6 x 10^−4^; MMR-: *t_anc_* = 16.10, *P_anc_* = 5.4 x 10^−10^; *t_1-day_* = 6.16, *P_1-day_* = 4.5 x 10^−5^; *t_100-day_* = 2.22, *P_100-day_* = 0.039). Photos of **(c)** 1-day (P108), **(d)** 10-day (P408), and **(e)** 100-day (P508) populations co-cultured with the WT ancestor on TA agar and incubated at 37°C for 24 h to show variation in colony size. Colonies from evolved populations appear red while WT ancestor colonies appear beige. Inset in (**d)** provides a magnified look at ancestor and 10-day colonies; the black arrow points to a 10-day population colony, highlighting the evolved extreme reduction in colony size. (**f**) Presence/absence of biofilm in evolved cultures after 24 h. Surface biofilm can be observed in daily and 10-day populations, but is absent in 100-day populations and the ancestor. (**g**) Average density of biofilm produced by evolved populations and ancestors in 96-well plate assay after 24 hours of growth, quantified via absorbance at 550 nm. Populations evolving to 10-day resource limitation cycles produce significantly more biofilm (WT: *t_anc_* = 10.63, *P_anc_*= 7.0 x 10^−10^; *t_1-day_* = 8.97, *P_1-day_* = 5.0 x 10^−9^; *t_100-day_* = 10.25, *P_100-day_* = 7.0 x 10^−10^; MMR-: *t_anc_* = 15.37, *P_anc_* = 1.6 x 10^−15^; *t_1-day_* = 8.22, *P_1-day_* = 7.7 x 10^−11^; *t_100-day_* = 14.21, *P_100-day_* = 7.5 x 10^−15^).

Previously, we observed that early biofilm formation was one of the major growth phenotypes that emerges in our culture conditions ^30^. Examination of culture tubes after 24 h revealed that 1-day and 10-day populations produce visible biofilms at the surface-air interface, while ancestral and 100-day populations generally do not (**Fig. 1f**). Further, quantification of biofilm formation reveals that 10-day populations produce significantly thicker biofilms than ancestor, 1-day, and 100-day populations (**Fig. 1g**). This languid early growth combined with increased capacity for biofilm formation in 10-day populations illustrates a pattern of non-monotonic evolution for growth phenotypes in response to chronic resource limitation.

Next, as the mutation rate has been reported to increase during acute starvation, we performed fluctuation tests for rifampicin resistance ^31^ on clones isolated from evolved populations to check for evolved changes in the mutation rate. For populations originating from WT backgrounds, all 12 clones isolated from 10-day resource-limitation cycles exhibited significantly higher rates of rifampicin resistance than the WT ancestor (**Fig. 2a**). This evolved widespread increase in mutation rate is unique to populations from 10-day resource-limitation cycles, as significant increases were only observed for three clones isolated from 1-day populations and none from 100-day populations. In MMR-backgrounds, mutation-rate patterns were similar, as six out of twelve clones isolated from 10-day resource-limitation cycles evolved higher rifampicin resistance rates than the MMR-ancestor, while only two and three tested clones show such significance from 1-day and 100-day populations, respectively (**Fig. 2a**). Thus, a trend of non-monotonic evolutionary responses is also observable in mutation rates -- cultivation in the intermediate, 10-day resource-limitation cycle often results in the evolution of an increased mutation rate, while cultivation at the other extremes commonly results in relatively little change in mutation rate.

**Figure 2.**
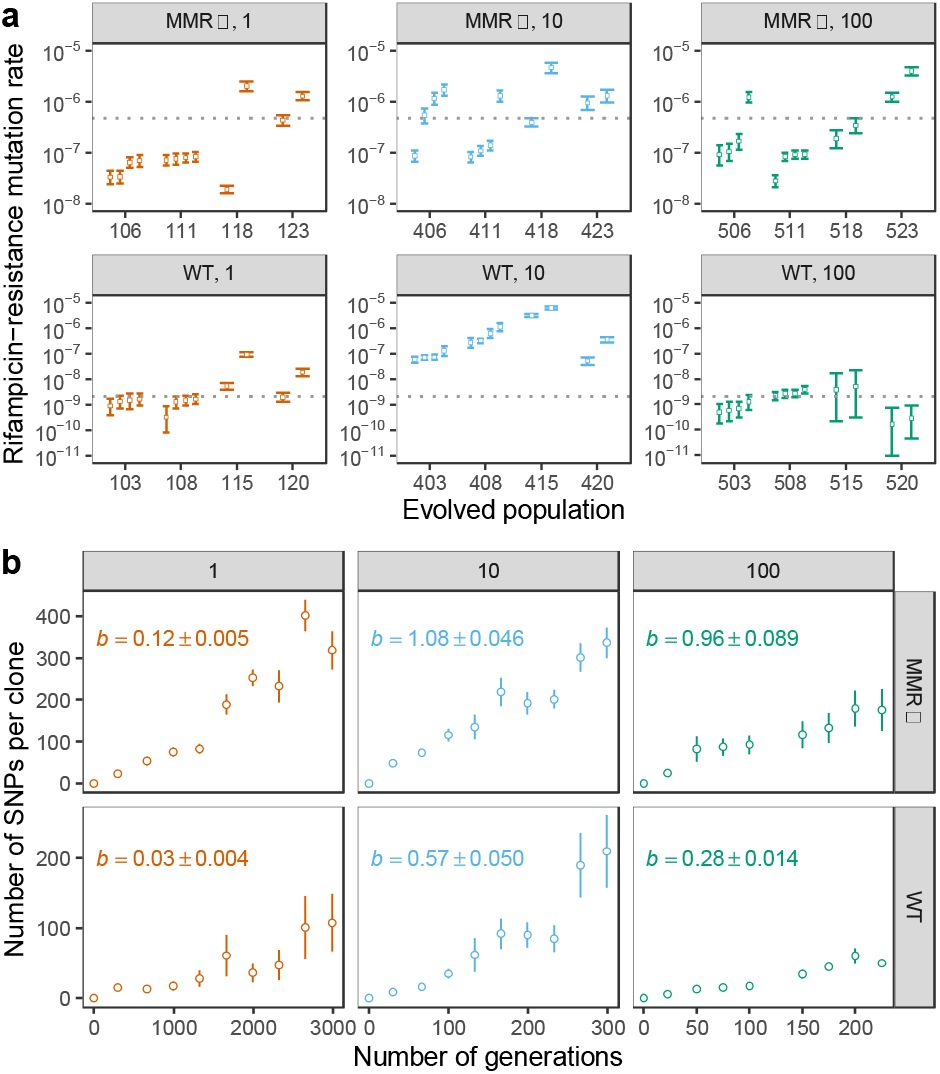
Evolutionary dynamics of evolved populations. **(a)** Each panel shows mutation rates of populations after 900 days of experimental evolution in different combinations of resource-limitation cycle length (1, 10, or 100-days) and genetic background (MMR-for mismatch repair defective or WT for wild-type). In each combination, four evolved replicate lines were tested, and two or four clones per evolved line were isolated and measured. As mutation rates are estimated via maximum-likelihood, the open circle and error bar represent the mean and the 95% confidence interval for each clone. The grey dashed line represents the mutation-rate measurement of the corresponding ancestor. The transparent coloured lines represent the mean mutation-rate measurement of each combination. For both WT and MMR-backgrounds, populations evolving to 10-day resource limitation cycles exhibit greater mutation rates than the other treatments (WT: *P* < 10^−4^; MMR-: 0.081; permutation test). **(b)** Each panel shows the mean rate of genomic evolution among the parallel experimental populations for each resource-limitation cycle length (1, 10, or 100 days) in MMR- (mismatch repair defective) or WT (wild-type) genetic backgrounds. At each time point, the circle and the bar represent the mean and the standard error. From left to right represent the sequencing profiles sampled at days 0, 90, 200, 300, 400, 500, 600, 700, 800, and 900. Note that the data for populations with 100-day resource-limitation cycles are missing at day 500. The line represents the linear regression against the number of generations. The estimated slope (*b*) and corresponding standard error are also shown in each panel.

As the effects of individual mutations can vary, it is possible for phenotypes and genotypes to evolve at different rates and produce different evolutionary response patterns ^32^. To determine if the rate of genomic evolution also exhibits a similar non-monotonic pattern as observed in growth dynamics and mutation rate, we assessed genomic-sequence divergence throughout 900 days of evolution. Using high-throughput population-genomic data collected every 100 days (mean coverage > 100×, **Supplemental Table 1**), genomic divergence was estimated by summing the derived-allele frequencies for all single nucleotide polymorphisms (SNPs) at each time point. The rate of genomic evolution is then defined as the slope of the linear regression of summed derived-allele frequencies against the estimated number of generations for each resource-limitation treatment. Consistent with the magnitudes of change observed in growth dynamics and mutation rate, 10-day populations also experienced the highest genomic evolution rates (**Fig. 2b**). This pattern is true regardless of initial genetic background, and alternative measures of genomic divergence do not qualitatively affect the results (**Supplemental Fig. 1a-d**). In addition to the genomic evolution rate, comparison of the fixation probabilities for detectable nonsynonymous, intergenic, and synonymous mutations, suggests that positive selection is not significantly stronger in 10-day populations than the other evolutionary conditions. Instead, it is more likely that the increased mutation rates that evolve in response to 10-day resource-limitation cycles are the greatest contributing factor to the observed increases in the genomic evolution rate (**Supplemental Fig. 1e-g**).

Although genomic and phenotypic evolutionary responses to resource availability diverge from monotonically increasing or decreasing functions, the interaction of these responses may result in different patterns for intraspecific diversity and community structure. For 1-day cycles of resource limitation, our culture environment is known to support subpopulation structure ^30^. However, how increases in the length of resource-limitation cycles differentially affects the ability to create and maintain subpopulation structure remains unknown. We used a clade-aware hidden Markov chain (caHMM), which assumes the coexistence of two clades (major and minor), to infer whether each mutation belongs to the basal clade (i.e., mutations that sweep through the entire population before the establishment of subpopulations), major clade, or minor clade. This approach also infers if a mutation remains polymorphic or reaches fixation within its assigned clade at each sample time point ^33^.

Application of caHMM revealed that resource-limitation cycle length affects the persistence of subpopulation structure, as the durations of time that major and minor clade are inferred to coexist (**Fig. 3a**) are significantly shorter in 100-day populations than in 1-day and 10-day populations (**Fig. 3b**). Consistent with this observation, the proportion of fixed mutations assigned to the basal clade in a population is negatively correlated with the duration of subpopulation structure persistence (**Supplemental Fig. 2a**), being greatest in 100-day populations (**Supplemental Fig. 2b**). Plotting the temporal distribution of derived-allele frequencies reveals that mutations with intermediate frequencies (0.3-0.7) are less abundant in 100-day populations (**Supplemental Fig. 2c**), again consistent with a lower tendency to form and further maintain subpopulation structure.

**Figure 3.**
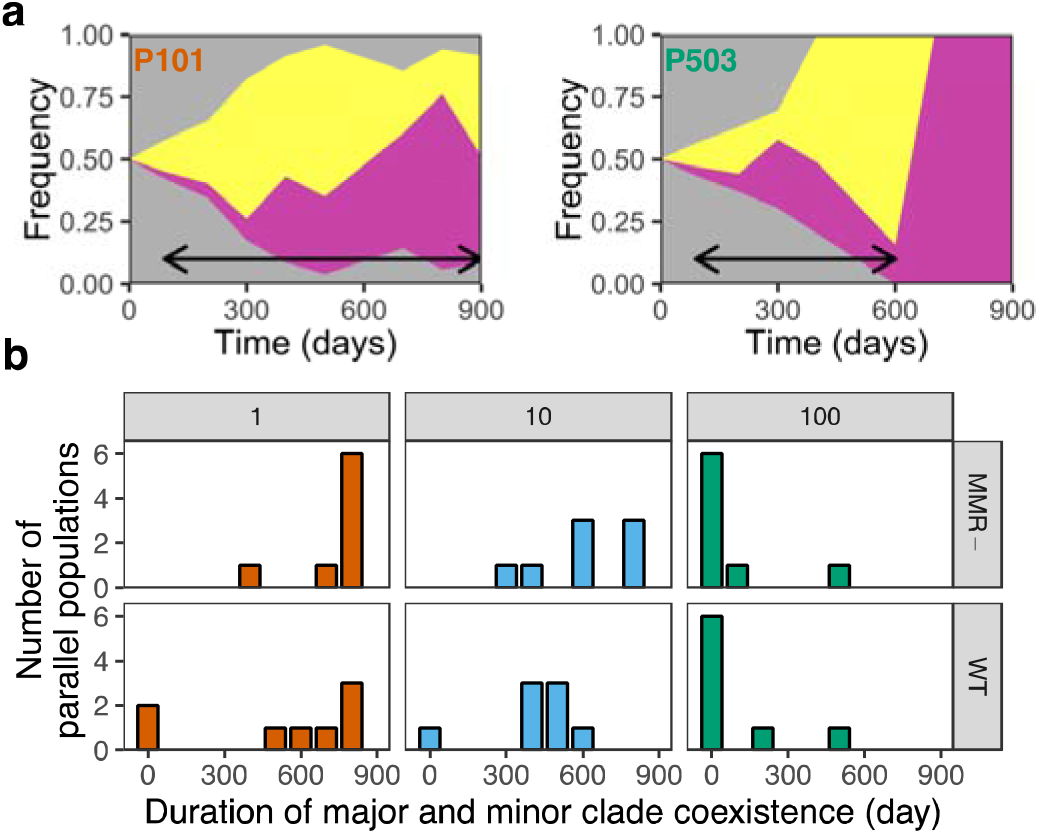
Ecological dynamics in experimental evolution. Muller plots of major (yellow) and minor (pink) clade abundance for (**a**) Population 101 (1-day, WT) and Population 503 (100-day, WT). Black arrows denote the maximum duration that two clades are observed to co-exist based on the presence of associated SNPs identified with metagenomic sequencing. (**b**) The distribution of durations of coexistence of major and minor clades, or subpopulation structure, as determined by the caHMM among the parallel experimental populations for each resource-limitation cycle/genetic background combination – 1-day (orange), 10-day (blue) and, 100-day (green). The duration of coexistence for multiple clades in a population decreases as the duration of resource limitation cycles increase (*P* = 6.1 x 10^−6^, Scheirer-Ray-Hare test), with populations evolving to 100-day resource limitation cycles exhibiting the least subpopulation structure (1-day v. 100-day: *P* = 3.1 x 10^−5^; 10-day v. 100-day: *P* = 4.2 x 10^−5^, Mann-Whitney *U* test).

To ensure that a lower tendency to support subpopulation structure is unlikely to be an artefact of insufficient evolutionary time, we normalized our comparisons by the number of derived mutations observed in each evolutionary treatment. For example, 1-day populations have experienced the same amount of genomic evolution after 300-days of culture as 100-day populations have experienced after 500-days of culture (calculated based on **Fig. 2b**). However, by 300-days, fixation of mutations in the basal clade slowed and subpopulation structure stabilized in 13 of 16 daily-transferred populations. Conversely in 100-day populations, although the same amount of molecular evolution occurred by 500-days, subpopulation structure only appeared to stabilize in 2 of 16 populations. Thus, evolutionary outcomes relating to the maintenance of subpopulation structure in response to resource limitation can be described by a decreasing monotonic function – such that, as the duration between resource replenishment increases the ability to sustain subpopulation structure decreases. This also reveals that the interaction of forces shaping the previously mentioned phenotypic and genotypic evolution in response to resource-limitation may result in discordant patterns with the evolution of subpopulation structure.

Lastly, to distinguish which mutations are likely positively selected and adaptive from mutations that are likely promoted to fixation due to genetic draft ^34^, we focused on the nonsynonymous mutations which were inferred to be fixed within either the basal, major, or minor clades of the evolving populations. Here, two metrics were calculated which indicate the degree of parallel evolution within a treatment combination and further identify genes which are likely selective targets due to an overrepresentation of fixed nonsynonymous mutations: sum of *G*-scores ^35^ and Bray–Curtis similarity ^15^. Moreover, if the fixed non-synonymous mutations are concentrated in a smaller subset of genes within a resource-limitation/genetic background combination then the resulting values for these two metrics will further increase. For all treatment combinations, both the sum of *G*-scores and the Bray-Curtis similarity values were significantly greater than expected based on a simulated null distribution (**Fig. 4a-b**). Thus, revealing a significant amount of parallel evolution within treatments and suggesting that positive selection is likely shaping these populations. Interestingly, the statistical significance of both metrics for 10-day and 100-day populations is also much greater than for 1-day populations, suggesting a smaller pool of adaptive targets and that ability of the 10-day and 100-day populations to explore the adaptive landscape is more limited, possibly due to their environmental harshness.

**Figure 4.**
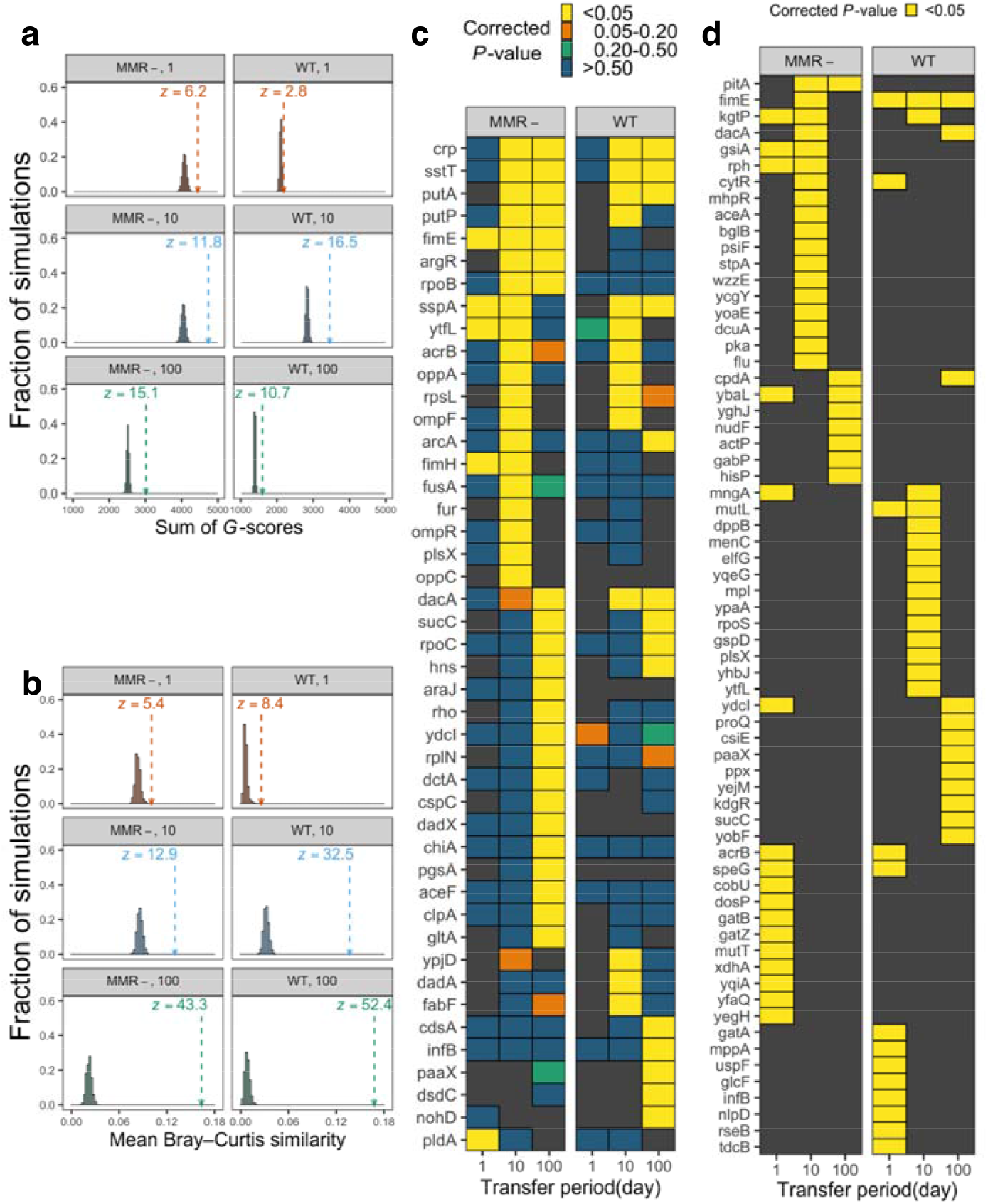
Evolutionary parallelism as evidence for positive selection. Within each resource-limitation cycle/genetic background combination, the sum of G-scores was computed - a goodness of fit test that can indicate the extent of parallel evolution and identify genes that are likely selective targets. **(a)** The arrow and vertical dashed line show the observed sum of *G*-scores for each treatment combination: 1-day (orange), 10-day (blue), and 100-day (green). The histogram represents the null distribution of evolutionary parallelism as determined by 20,000 simulated sums of *G*-scores, where the position of mutations were randomized throughout the genome. The significance of the observed sums can be evaluated by *z*-scores which quantify the extent the observed value deviates from the null distribution (*z* > 1.65 for one-tailed *P* < 0.05). **(b)** Similarly, the arrow and vertical dashed line show the observed mean of Bray–Curtis similarity across the pairs of evolved populations evolved in a resource-limitation cycle/genetic background combination. The histogram represents the null distribution of evolutionary parallelism as determined by 1,000 simulated means of Bray–Curtis similarity, where the position of mutations were randomized throughout the genome. The significance of the observed sums can again be evaluated by *z*-scores (*z* > 1.65 for one-tailed *P* < 0.05). **(c)** List of genes with at least one nonsynonymous mutation that significantly contribute to the increased sum of G-scores and are likely under positive selection. Significance levels (simulated *P*-values with Bonferroni correction) are shown by the different non-black colours of tiles. Genes with no such hits in a particular resource-limitation cycle/genetic background combination are shown by black tiles. **(d)** List of genes significantly overrepresented for structural mutations (IS-element insertions and indels) that are likely under positive selection. Yellow tiles highlight genes observed in at least two populations and yielding simulated *P*-values < 0.05 after Bonferroni correction.

To more deeply investigate the observed parallelism in likely adaptive mutations, we conducted gene ontology analyses focusing on genes that were determined to significantly contribute to the increased sum of *G*-scores in each treatment combination (**Fig. 4c; Supplemental Table 2**). We additionally investigated genes which were overrepresented for disruptive ‘structural’ mutations such as indels or IS-element insertions (**Fig. 4d; Supplemental Table 3**). Furthermore, an additional set of 115 nonsynonymous mutations which were parallel at the nucleotide level – that is, these mutations independently arose and are responsible for identical amino-acid substitutions in at least two experimental populations within a treatment combination (**Supplemental Fig. 3a-b; Supplemental Table 4**). Results of the gene ontology analyses revealed that genes associated with DNA repair (GO:0006281) or replication (GO:0006260) more commonly experienced mutations in 10-day populations than in 1- or 100-day populations (**Supplemental Table 5**), consistent with the evolution of increased mutation-rates in 10-day populations. However, some parallelism was observed which may indicate a general response to resource limitation as genes related to global gene expression (regulation, termination, and fidelity of transcription and translation), resource import, and metabolism (**Supplemental Table 6-9**) were enriched for mutations in both 10- and 100-day populations.

Since populations evolving in response to 10-day and 100-day resource-limitation cycles exhibited parallelism in biological processes that are enriched for likely adaptive mutations but differed so greatly in their resulting phenotypes, we hypothesized that treatment-specific differences must exist in the individual genes or amino acids experiencing mutations. Here, notable differences in parallel amino-acid substitutions occurred in the stringent response regulator *crp* (Y42C, 10-day populations; E56K, 100-day populations), the transcription termination factor *rho* (R109H, 100-day populations), RNA polymerase subunit *rpoB* (D915G, 10-day populations; E859K, 100-day populations), and the translation elongation/ribosome recycling factor, *fusA* (R437C, I654N, 10-day populations) (**Supplemental Table 4**). Although these genes are all involved in processes that have broad effects on gene expression, it is plausible that the mutation of these genes may result in widely-different phenotypic effects. Particularly, in the case of *crp*, as the mutations that arise in 10-day and 100-day populations are located in or proximal to the RNA polymerase-interacting domain. Here, mutation variants have been previously characterized to have significantly different effects on *crp*’s interaction with RNA polymerase and the activation of promoters ^36,37^. Thus, further studies are needed to understand how the alteration of genes that are responsible for these basic molecular processes is beneficial to populations experiencing distinct levels of repeated starvation – both in terms of differential expression and the resulting phenotypes.

## Discussion

Several conclusions can be drawn from the reported results. First, we observed that the pattern of evolutionary responses for many molecular and phenotypic traits do not follow simple increasing or decreasing functions across an environmental gradient of resource limitation, i.e., they are non-monotonic. This is at least qualitatively consistent with previous theoretical and experimental studies of fluctuating environments, suggesting that periodic stress results in higher phenotypic variability than longer, more sustained stress – increasing their potential to evolve novel adaptations ^38,39^. In addition to changes in biofilm density, growth rate, and colony size, we also notably see this phenomenon extend to the evolution of increased mutation rates, which are often induced in stressful environments ^40^. Consistent with our results, population-genetic models have suggested that higher mutation rates can evolve in fluctuating environments and increase in prevalence due to second-order selection ^41–43^. In these cases, mutators are expected to be the most prevalent in environments where conditions fluctuate at intermediate frequencies ^44^. Thus, taking into account these observed non-monotonic patterns, our results highlight the risk of assuming that expected evolutionary outcomes in response to intermediate levels of resource limitation can be inferred from observations in more extreme environments.

Second, evolutionary patterns can be discordant between phenotypic, genomic, and intraspecific community-structure outcomes. While we observed a non-monotonic pattern in the evolution of growth dynamics, mutation rate, and genome-evolution rate, persistence in subpopulation structure instead declined monotonically. Specifically, we observed that the duration during which multiple clades coexisted within a population decreased with longer resource-limitation cycles. This echoes the findings that resource-deficient environments promote detachment of biofilm formers ^45^ or metastasis of cancer cells ^46^, both of which increase population homogeneity. A pattern of decreasing subpopulation structure may arise due to a reduced pool of beneficial mutations combined with increased population bottlenecks in more resource-limited populations, limiting the ability to explore the adaptive landscape ^47^, In effect, this would result in a decreased potential for clonal interference and adaptive diversification, two phenomena that would otherwise cause the appearance of coexisting clades.

Lastly, albeit surprisingly, repeated cycles of resource limitation do not appear to promote the fixation of mutations affecting genes commonly associated with the growth advantage in stationary phase (GASP) phenotype, such as *rpoS* ^21,48^ and *Irp*^22^. In fact, other global regulators and genes with broad effects on transcript and protein expression (*crp*, *rho, rpoB*, and *fusA*) are instead targets for likely adaptive mutations. Further investigation is needed to determine if the absence of known GASP-associated mutations is due to strain-specific/culture environment differences or because these mutations are transient, with benefits that are no longer realized after repeated exposure to extreme starvation.

## Materials and Methods

### Construction of ancestral strains and experimental evolution

Experimental populations originated from one of four ancestral strains. Populations denoted as WT are descendants of PMF2, a prototrophic derivative of *E. coli* K-12 str. MG1655, while populations denoted as MMR-are descendants of PMF5 (PMF2, Δ*mutL*) provided by the Foster Lab ^29^. To introduce a neutral marker that would allow for the periodic screening of contamination between populations, lambda red recombineering was used to introduce a 3513 bp deletion removing the *araBAD* operon from one-half of both the WT and MMR-starting strains. This deletion was confirmed by the appearance of red colonies after plating on TA agar (1% Arabinose, 1%Tryptone, 0.5% NaCl, 0.1% Yeast Extract, 0.005% TTC (Sigma T8877)) and next-generation sequencing.

Long-term cultures were established by inoculating 10 mL of LB-Miller broth (Dot Scientific) with a single-isolated progenitor colony cultivated overnight at 37 °C on LB agar plates. Resulting cultures were maintained in 16- × 100 mm glass culture tubes containing 10 mL LB broth shaking at 175 rpm at 37°C. Selection conditions involved subjecting cultures to varying degrees of repeated starvation in which 1 mL of thoroughly-vortexed culture was transferred into 10 mL of fresh LB broth every day, every 10-days, or every 100-days. To maintain an historical record of the populations and to revive populations in the case of contamination or extinction, 1 mL of culture was collected pretransfer every 30 days from daily and 10-day populations, frozen with liquid nitrogen, and stored at −80°C. In order to not disrupt any potential ecological structure in 100-day populations, copies of these cultures were only collected pretransfer every 100-days. In addition, 100-day populations were maintained in triplicate (1 main culture and 2 backup cultures) with only the main culture being used to seed new triplicate cultures every 100 days. In the case of a whole population extinction, one backup culture would replace the main culture and the experiment would continue. This practice was only necessary during the first 200-days as past this time point, no additional whole-population extinction events occurred. Populations were screened for cross-contamination on days 90, 200, 300, 400, 500, 600, 700, 800, and 900 by streaking on MacConkey Agar (BD Difco) supplemented with 0.4% Arabinose.

### Evaluation of growth phenotypes

Growth curves were measured using an Epoch2 Microplate Spectrophotometer (Biolog). To prepare cultures for growth rate measurement, 900-day populations and day-0 progenitor strains were revived frozen −80°C stock by transferring a scrape of ice to 16- x 100 mm culture tubes containing 10 mL of LB broth. Cultures were incubated overnight at 37°C and shaking at 175 rpm. Following overnight culture, 1.5 μL of each revived population or progenitor were transferred into a well in a 96-well flat bottom culture plate (Corning/Falcon, 353072) containing 150 μL of LB broth. Evolved populations were measured as an average of eight replicates and progenitor strains were measured as an average of 16 replicates, with an additional eight wells dedicated to negative controls consisting solely of LB broth. After inoculation, lids were placed on 96-well plates and were sealed with parafilm before placement in the spectrophotometer. Throughout the duration of growth curve measurement, cultures were maintained at 37°C, with double orbital shaking at 548 cpm. Growth measurements were taken every 15m over the course of 16h at an OD of 600 nm. Maximum growth rate (*r*_max_) was calculated for all measurable replicates (WT: *n_ans_* = 32, *n_1-day_* = 16, *n_10-day_* = 16, *n_100-day_* = 16; MMR: *n_ans_* = 29, *n_1-day_* = 16, *n_10-day_* = 12, *n_100-day_* =16) using the Gen5 software (Biotek) as the maximum slope between two timepoints and time to *r*_max_ (*T*_max_) was recorded as the midpoint between *r*_max_ determining timepoints. For daily lines experiencing diauxic growth, calculations of *r*_max_ and *T*_max_ were restricted to the first logistic growth phase.

To measure colony size, revived 900-day populations and day-0 progenitor strains were grown overnight at 37°C and shaking at 175 rpm in 16- x 100 mm culture tubes containing 10 mL of LB broth. Following overnight incubation, serial dilutions of 100 μL of culture in 900 μL of phosphate buffered saline were made down to dilutions of 10^−5^, before plating 100 μL of diluted culture on to TA agar (1.6% agar, 1% arabinose, 1% tryptone, 0.5% NaCl, 0.1% yeast extract, 0.0005% TTC) and incubating for 24h at 37°C. TA plates were then photographed with a 20 mm scale in frame using an iPhone XS. Photos were then analysed with ImageJ (NIH, https://imagej.nih.gov/ij/), by first setting the pixel to distance ratio using the in-frame scale before flatting the photo to 8-bit black and white, setting the threshold to highlight the colonies, and using the ‘measure particles’ tool. Measurements for six arbitrary colonies per analysed population were recorded.

Presence and absence of biofilm at the surface-air interface was determined by reviving 900-day populations and day-0 progenitor strains in 16- x 100 mm culture tubes containing 10 mL of LB broth overnight at 37°C and shaking at 175 rpm. Following incubation, culture tubes were visually examined for the presence of biofilm attached to the culture tube at the surface-air interface. Quantification of biofilm formation in evolved populations was conducted using a microtiter plate biofilm assay ^49^. In a non-treated 96-well plate (Corning/Falcon, 351172), 15 μL of overnight culture was inoculated into 150 μL of LB broth. Each population was measured for a total of eight replicates with each 96-well plate containing two columns (16-wells) of *araBAD*-WT ancestor as a control and at least two columns (16-wells) of blanks. Inoculated 96-well plates were incubated for 24 h at 37°C. Following incubation, plates were washed with 1x PBS, stained with 0.1% crystal violet for 10 m (200 μL per well), washed again with 1x PBS, and left to dry overnight. Crystal violet bound to biofilms was then solubilized with 30% acetate for 15 m (200 μL per well), transferred to a fresh 96-well plate, and absorbance was quantified with a using an Epoch2 Microplate Spectrophotometer at 550 nm.

### DNA isolation and high-throughput sequencing

High resolution population tracking was conducted by collecting 1 mL of culture at day 90, 200, 300, 400, 500, 600, 700, 800, and 900 of experimental evolution. DNA was extracted using the DNeasy UltraClean Microbial Kit (Qiagen 12224; formerly MO BIO UltraClean Microbial DNA Kit) and submitted to either the Hubbard Center for Genomic Analysis at the University of New Hampshire, the Center for Genomics and Bioinformatics at Indiana University, or the CLAS Genomics Facility at Arizona State University for library preparation and sequencing. Libraries were generated with the Nextera DNA Library Preparation Kit (Illumina, FC-121-1030) following an augmented protocol for optimization of reagent use^50^ before being pooled and sequenced as paired-end reads to a target depth of 100x on an Illumina HiSeq 2500 (UNH) or an Illumina NextSeq 500 (Indiana; ASU).

### Sequencing analysis

Sequencing analysis was performed on the Mason and Carbonate high-performance computing clusters at Indiana University. Sequencing reads were quality controlled using Cutadapt v.1.9.1^51^ to remove residual adapters and trim low quality sequences. QCed sequences were then mapped to the *Escherichia coli* K-12 substr. MG1655 reference genome (NC_000913.3) and all mutations and their frequencies were identified using Breseq v.0.30.2 with the predict-polymorphisms parameter setting^52^. All mutations that were identified to previously exist in ancestral lines were discarded from the following analysis. Further, our analysis only included the samples which passed the following additional four quality checks: 1) we required mean sequencing depths > 10; 2) any WT sample identified to contain the 1,830 bp deletion in *mutL* from the PMF5 progenitor strain was discarded; 3) regions lacking sequencing coverage (i.e. depth = 0) must be smaller than 5% of the genome; and 4) the sequencing result should reflect the correct genetic background in terms of *ara* markers, including a nonsynonymous SNP at position 66528, an intergenic SNP at position 70289, and a multiple base substitution mutation (SUB) at position 66533. For an *ara*+ line, we required either of two SNPs showing DAF < 0.2. For an *ara*− line, we required either of two SNPs showing DAF > 0.8 or the SUB is detected. In the end, 395 genomic profiles passed QC and were included in the following analysis (**Supplemental Table 1**).

In order to remove the mutations that originated from the starting clone before experimental evolution, we discarded any mutations with a DAF = 100% at one time point for at least 11 experimental populations with the same genetic background from the analysis. In addition, as the highly repetitive sequences in *rsx* genes are known to cause errors in SNP calling^53^, they were also removed from the analysis.

### Guaranteed generations

The calculation of guaranteed number of generations is based on the following two observations. First, pre-transfer population density in the culture tube was measured at day 200, 300, 400, 500, 600, 700, 800, 900, by serial dilution in phosphate-buffered saline (PBS) before plating on LB agar for CFU counts and revealed that the population density does not significantly change throughout experimental evolution (**Supplemental Fig. 4**). This suggests the carrying capacity of experimental populations is recovered within a resource-limitation cycle after 1:10 dilution and the populations must have experienced at least 3.3 (log_2_10) generations between resource-limitation cycles. Therefore, the first estimate of guaranteed generations (*g*_1_) per day equals 3.3, 0.33, and 0.033 when the resource-limitation cycle is one day, 10 days, and 100 days, respectively. Second, we found that between resource-limitation cycles novel mutations arise and are able to be fixed within one sequencing interval. Therefore, given the carrying capacity *K*, the population should at minimum experience log_2_*K* cell divisions in 100 days. Using estimated *K* = 1.21 x 10^10^, 2.13 x 10^8^, and 3.71 x 10^7^ (**Supplemental Fig. 4**), the second estimate of guaranteed generations (*g*_2_) per day equals 0.30, 0.28, and 0.25 when the resource-limitation cycle is one day, 10 days, and 100 days, respectively. The final number of guaranteed generations (*g*) for each resource-limitation cycle is then determined as the larger value between the corresponding *g*_1_ and *g*_2_.

Given that *K* was only estimated pre-transfer, when the feeding cycle is long it is possible that populations have reached to a density *K*\* higher than *K* and decreased back to *K* within one feeding cycle. If this is true, the first estimated guaranteed generations (*g*_1_’) per day should be [log_2_(*K*\*/*K*) + 3.3]/RC, where RC is the length of the resource-limitation cycle. While *g*_1_’ is likely to be larger than *g*, the observed fold-changes in the genomic evolutionary rates between in longer resource-limitation cycles cannot be fully explained by this possibility. For example, if the 9.4-fold change in 10-day lines compared to daily lines were entirely due to this possible underestimation, *g*_1_’ needs to be 9.4-fold higher than *g*_1_, which requires an implausible *K*\* = 4.7 x 10^16^. Similarly, if the 8.8-fold changes in 100-day lines compared to daily lines were entirely due to this possible underestimation, *g*_1_’ needs to be 8.8-fold higher than *g*, which also requires an implausible *K*\* = 7.2 x 10^64^. Therefore, even if we had tracked the population density throughout the entire experimental evolution, it is likely that we still could not find substantial effects of changing population densities on our observations.

### Rate of genomic evolution

We quantified the level of genomic divergence for each experimental population at each time point by summing all DAFs of detected mutations. Then we calculated the mean genomic divergence across all eligible experimental populations in each resource-limitation cycle/genetic background combination (**Supplemental Table 1**). When estimating the rate of genomic evolution, the linear regression was performed by the function “lm” in R with formula “mean genomic divergence ~ guaranteed generations + 0” (linear model), which enforces the y-intercept as 0.

To explore the possibility of nonlinear relationship between the genomic divergence and time, we also performed regression using the formula “mean genomic divergence ~ guaranteed generations + square root of guaranteed generations + 0” (nonlinear model), which was previously proposed to catch the trend of diminishing returns^35^. However, the results of nested ANOVA test suggest no strong evidence supporting that the nonlinear model is substantially better than the linear model (**Supplemental Fig. 1d**).

### Fluctuation test and mutation rate estimation

To quantify the genetic mutation rate of our evolved populations and their ancestors, we performed fluctuation tests as described^22^ on clones isolated from each evolved population at 900 days. Briefly, fluctuation tests measure the rate of resistance to the antimicrobial rifampicin which is conferred by mutations to *rpoB*. For each resource-limitation cycle/genetic background combination, the four *ara*− populations were assayed. For the first two populations with smaller population numbers, four independent clones were picked; for the last two populations with larger population numbers; two independent clones were picked. For each of the WT or MMR-ancestor, two independent clones were used. For each clone, 40 replicate experiments were performed. Number of mutants as determined by CFU counts/mL were converted to an estimated mutation rate and a corresponding 95% confidence interval by the function “newton.LD” function in the R package “rSalvador”^54^ for each clone.

When performing the permutation test for 36 WT clones (12 from 1-day populations, 12 from 10-day populations, and 12 from 100-day populations), we randomly shuffled their labels of population in 10,000 times. After each shuffle, we counted the number of clones exhibiting 95% confidence interval with the label of 10-day populations. The final *P*-value was determined by the proportion of these numbers larger or equal to the genuinely observed number exhibiting 95% confidence interval with the label of 10-day populations. The permutation test for 36 MMR-clones was similarly implemented.

### Clade-aware hidden Markov chain analysis (caHMM) and longest coexistence time

To test coexistence time in each experimental population, we performed clade-aware hidden Markov model (caHMM) using a modified version of previously released code^33^. Our modifications included: 1) modifying the code in “calculate_clade_hmm_wrapper.py,” in order to continue the iteration of experimental populations, even when the major clade frequency of all time points for an experimental population cannot be estimated; 2) removing the original cut-off for minimum coexistence time, as that parameter is too long for our experiments; and 3) increasing the iteration number for hidden Markov chain from 5 to 50. When performing caHMM, we constructed an annotated time course of mutations using the original format. If a mutation is a structural mutation (DEL, INS, and MOB), the sequencing depth is determined by the mean sequencing depth across the mapped adjacent nucleotide (“coverage_plus” or “coverage_minus”) of associated junction candidates (JCs) in breseq annotated file. The sequencing depth of time 0 was set as 100.

Using the major clade frequency (*f*_M_) and minor clade frequency (*f*_m_) estimated by caHMM, we calculated the longest coexistence time as follows: if there is no time point showing 0.2 < *f*_M_ <0.8 and 0.2 < *f*_m_ <0.8, its longest length of coexistence is defined as zero; otherwise, we found the longest time intervals within which every time point shows 0.01 < *f*_M_ <0.99 or 0.01 < *f*_m_ <0.99. After calculating the longest length of coexistence for each population, to test whether length of resource-limitation cycle and genetic background affects the longest length of coexistence, we performed a Scheirer-Ray-Hare test, (nonparametric version of two-way ANOVA) using the function “scheirerRayHare” in the R package “rcompanion”^55^. The sample size for each resource-limitation cycle/genetic background combination is eight. The succeeding *post hoc* analysis is done by Mann-Whitney *U* test with the function “wilcox.test“ in R.

### Identification of candidate mutations for positive selection

For the populations where caHMM does not finish, we instead performed well-mixed hidden Markov chain (wmHMM) using the code “calculate_well_mixed_hmm_wrapper.py,” which also infers the fixation status of a mutation under a one-clade model. The single clade in wmHMM is defined as the basal clade. Candidate mutations for adaptation are then defined as mutations that satisfy either of the following two criteria: (1) the mutations recognized as fixed mutations in any of the basal, major, or minor clade in the output file from caHMM; (2) the mutations that show DAF > 0.5 in at least two time points.

### Calculation of G-scores

To quantify the parallelism of the set of candidate nonsynonymous mutations in a resource-limitation cycle/genetic background combination, we calculated *G*-score for each gene^35^. A larger *G*-score means more overrepresentation for a gene. To calculate *G*-scores, we first counted the observed number of candidate nonsynonymous mutations in gene *i* per resource-limitation cycle/genetic background combination (*O*_i_), and the expected number for gene *i* (*E*_i_) was calculated by *O*_tot_(*L*_i_/*L*_tot_), where *O*_tot_ = ∑_i_ *O*_i_, *L*_i_ is the number of nonsynonymous sites for gene *i*, and *L*_tot_ = ∑_i_*L*_i_. The *G*-score for gene *i* (*G*_i_) was then calculated by 2*O*_i_ln(*O*_i_ /*E*_i_) or defined as zero when *O*_i_ = 0. Note that 2*O*_i_ln(*O*_i_ /*E*_i_) could be smaller than zero when 0 < *O*_i_ < *E*_i_. This can lead to a paradox that *G*_i_ is smaller when a gene was hit by a few times compared to the situation that the same gene was not hit at all. Therefore, we also defined *G*_i_ = 0 when 2*O*_i_ln(*O*_i_ /*E*_i_) < 0 in the following analysis.

Because the null expectation of *G*-scores varies with total number of fixed nonsynonymous mutations (**Supplemental Fig. 3b**), we performed 20,000 simulations in each of which *O*_tot_ hits are randomly distributed among all *L*_tot_ sites across all genes in the genome. To evaluate the significance of the sum of *G*-scores (**Fig. 4a**), we calculated the corresponding *z* score by (the observed sum - mean of simulated sums) / (standard deviation of simulated sums).

### Calculation of mean Bray-Curtis similarity

To quantify the parallelism of the set of candidate nonsynonymous mutations, we also calculate the mean Bray-Curtis similarity across all pairs of experimental populations for a resource-limitation cycle/genetic background combination. Specifically, for a pair of populations *j* and *k*, their Bray-Curtis similarity is calculated by

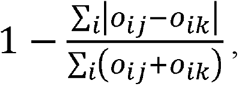

where *o*_ij_ and *o*_ik_ is the observed number of candidate nonsynonymous mutations in gene *i* for population *j* and *k*, respectively.

We also performed 1,000 simulations to acquire its null distribution for each resource-limitation cycle/genetic background combination. In each simulation, we randomly sample the nonsynonymous sites up to the number of observed candidate nonsynonymous mutations for each population and calculated mean Bray-Curtis similarity as described above. After acquiring the null distribution, to evaluate the significance of the mean Bray-Curtis similarity (**Fig. 4b**), we calculated the corresponding z score by (the observed value - mean of simulated values) / (standard deviation of simulated values).

### Overrepresentation of the genes affected by nonsynonymous mutations

To evaluate the significance of *G*-score for gene *i*, we directly compared the *G*_i_ to the distribution of 20,000 simulated *G*_i_. The *P*-value was determined by the proportion of simulated *G*_i_ larger or equal to the observed *G*_i_. The multiple test correction was performed by multiply each gene’s *P*-value by the number of genes with at least one hit by the set of fixed nonsynonymous mutations (Bonferroni correction). Only the genes with Bonferroni corrected *P*-value < 0.05 are called significant.

### Enrichment test of GO terms and KEGG pathways

Using this set of significant genes for each resource-limitation cycle/genetic background combination, we performed enrichment tests of gene ontology terms and KEGG pathways using the function “enrichGO” and “enrichKEGG”, the organismal database org.EcK12.eg.db, and a *q*-value cut-off = 0.05 in R package “DOSE”^56^.

### Overrepresentation of the genes affected by structural mutations

To test whether the set of genes that were affected by structural mutations (indels and IS-element insertions) in a resource-limitation cycle/genetic background combination were influenced by positive selection, we also calculated a *G*-score for each gene^35^. To calculate *G*-scores, we first counted the observed number of populations with any structural mutations in gene *i* per resource-limitation cycle/genetic background combination (*O*_i_), and the expected number for gene *i* (*E*_i_) was calculated by *O*_tot_(*L*_i_/*L*_tot_), where *O*_tot_ = ∑_i_ *O*_i_, *L*_i_ is the gene length for gene *i*, and *L*_tot_ = ∑_i_, *L*_i_. The *G*-score for gene *i* (*G*_i_) was then calculated by 2*O*_i_ln(*O*_i_ /*E*_i_), following the methods described in the above section.

We also performed 20,000 simulations and determined the Bonferroni corrected *P*-value for each gene *i* following the methods described in the above section. As a result, we found all the genes with *O*_i_ ≥ 2 show Bonferroni corrected *P*-value < 0.05.

### Data and Code Availability

Sequencing data generated during this study are available at NCBI’s Sequencing Read Archive: BioProject PRJNA532905 (Embargoed data accessible to reviewers: https://tinyurl.com/tsynzor). Code generated to analyze sequencing data and generate figures is available at https://github.com/LynchLab/ECEE_Starvation

## Acknowledgements

We thank B. Choi, T. Doak, D.A. Drummond, P. Foster, W. Guo, J. T. Lennon, H. Long, J. McKinlay, S. Snyder, E. Thorley, A. Urquidez, S. Vidrios, and S. Walls for their assistance and helpful comments. High Performance Computing Resources were provided and maintained by the National Center for Genome Analysis Support at Indiana University. This work was supported by Army Research Office Grants ARO65308-LS-MU and W911NF-14-1-0411 and National Institutes of Health Grant F32GM123703 and R35GM122566.

## Author Contributions

Conceptualization, M.G.B., W-C.H., and M.L.; Methodology, M.G.B., W-C.H., and S.F.M.; Investigation, M.G.B., W-C.H., S.F.M., G.F.B., and J.C.M.; Formal Analysis, M.G.B., and W-C.H.; Writing – Original Draft, M.G.B., and W-C.H.; Writing – Review & Editing, M.G.B., W-C.H., and M.L.; Visualization, M.G.B., and W-C.H.; Funding Acquisition, M.G.B. and M.L.

## Supplemental Information Titles and Legends

**Supplemental Figure 1.**
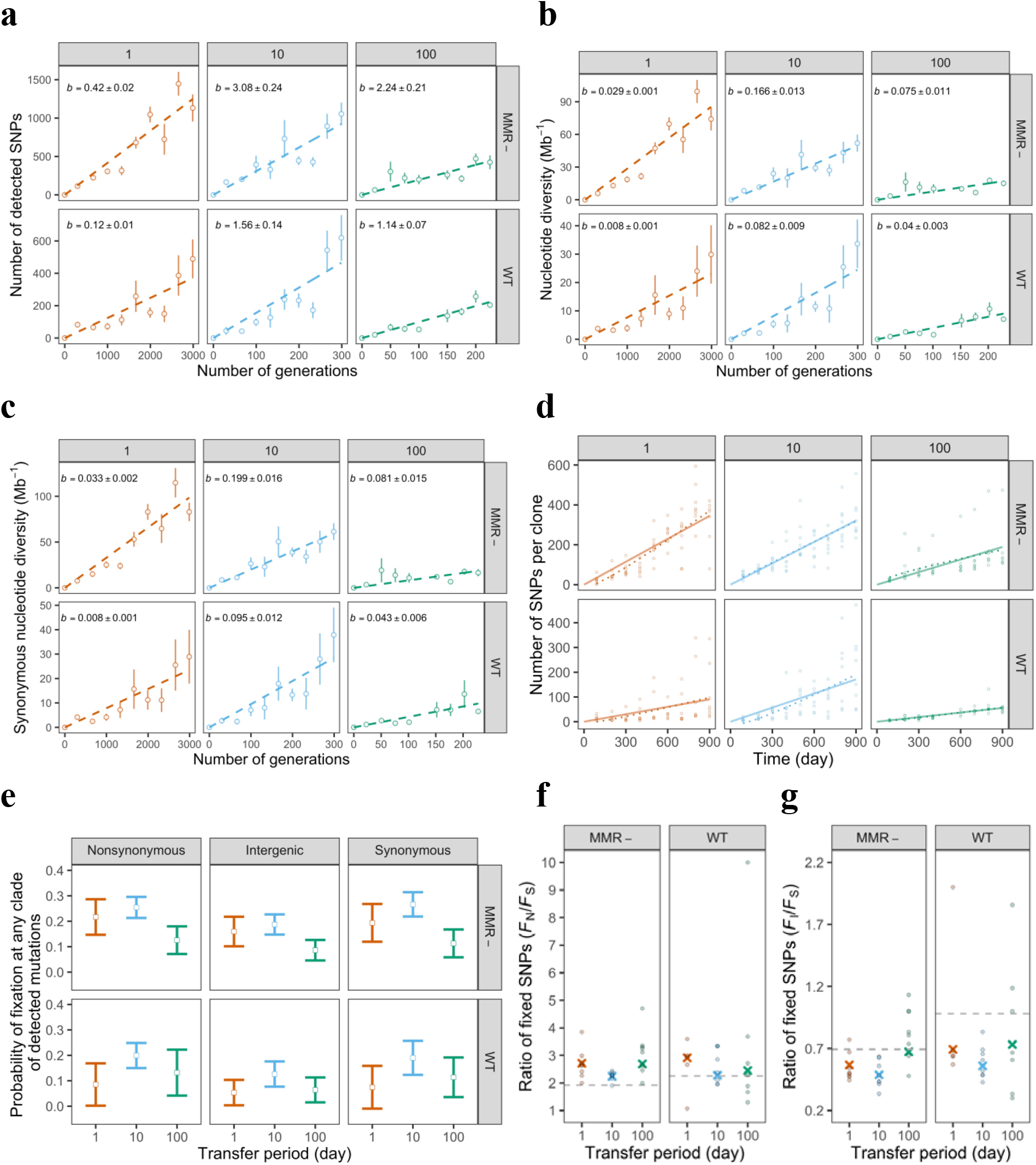
Genomic diversity changing over time and probability of fixation. Genomic diversity was measured by **(a)** number of SNPs per population, **(b)** nucleotide diversity, **(c)** synonymous nucleotide diversity, and **(d)** number of SNPs per clone, respectively. In **(a)**-**(c)**, the y-axis shows mean estimates among the parallel populations in a combination of one resource-limitation cycle (1 day, 10 days, or 100 days) and one genetic background (MMR-for mismatch repair defected, or WT for wild-type). Across the x-axis from left to right, circles represent the sequencing profiles sampled at day 90, 200, 300, 400, 500, 600, 700, 800, and 900 (the data of populations with resource-limitation cycle = 100days at day 500 are missing). At each time point, the circle and the bar represent the mean +/- standard error of mean. The dashed lines represent the linear regression lines of number of SNPs against guaranteed generations, and the estimated slopes (*b*) and corresponding standard errors are also shown. In **(d)**, the results of linear and nonlinear regression are compared. At each time point, a circle represents the number of SNPs per clone in a parallel population. The solid line represents the result of linear regression (*y* ~ 0 + *x*); The dotted line represents the result of nonlinear regression (*y* ~ 0 + *x* + *x*^0.5). Model comparison were performed by nested ANOVA: *P* = 0.028, 0.69, and 0.23 for resource-limitation cycle = 1, 10, and 100, respectively in MMR-background; *P* = 0.40, 0.063, and 0.40 for resource-limitation cycle = 1, 10, and 100, respectively in WT background. In **(e)**, similar probabilities of synonymous, nonsynonymous, and intergenic mutations fixed in any clade were found in populations with longer resource-limitation cycles. The dots and error bars represent mean probability and its 95% confidence interval (estimated by 1.96 times of standard errors) across parallel experimental populations. The fixed probably here is conditioning on the detectability of mutations. In **(f)**-**(g)**, populations with longer resource-limitation cycles do not show significant higher fixation rates of nonsynonymous mutations and intergenic mutations compared to the fixation rates of synonymous mutations. If positive selection is strong, one would expect to see that mutations with a greater potential to change protein functionality or levels (e.g. nonsynonymous mutations and intergenic mutations) are more likely to be fixed within a clade than mutations with lesser potential (e.g. synonymous mutations). Therefore, higher ratios of the number of nonsynonymous mutations fixed in any clade (*F*_N_) to the number of synonymous mutations fixed in any clade (*F*_S_), or higher ratios of the number of intergenic mutations fixed in any clade (*F*_I_) to *F*_S_, is expected. However, the above expectation was not observed. In **(f)**, the value represented by a cross is calculated by the summed *F*_N_ and *F*_S_ of all parallel experimental populations with a specific resource-limitation cycle length and genetic background. The value represented by a dot is calculated by the *F*_N_ and *F*_S_ in each parallel experimental population, and the dots with *F*_S_ = 0 are not shown. The dashed lines show the ratio of the number of nonsynonymous mutations to the number of synonymous mutations detected in mutation accumulation experiments. In **(g)**, the value represented by a cross is calculated by the summed *F*_I_ and *F*_S_ of all parallel experimental populations with a specific resource-limitation cycle length and genetic background. The value represented by a dot is calculated by the *F*_I_ and *F*_S_ in each parallel experimental population, and the dots with *F*_S_ = 0 are not shown. The dashed lines show the ratio of the number of intergenic mutations to the number of synonymous mutations detected in mutation accumulation experiments. Therefore, according to **(e)**-**(g)**, no evidence for significantly different levels of positive selection across resource-limitation cycles was found.

**Supplemental Figure 2.**
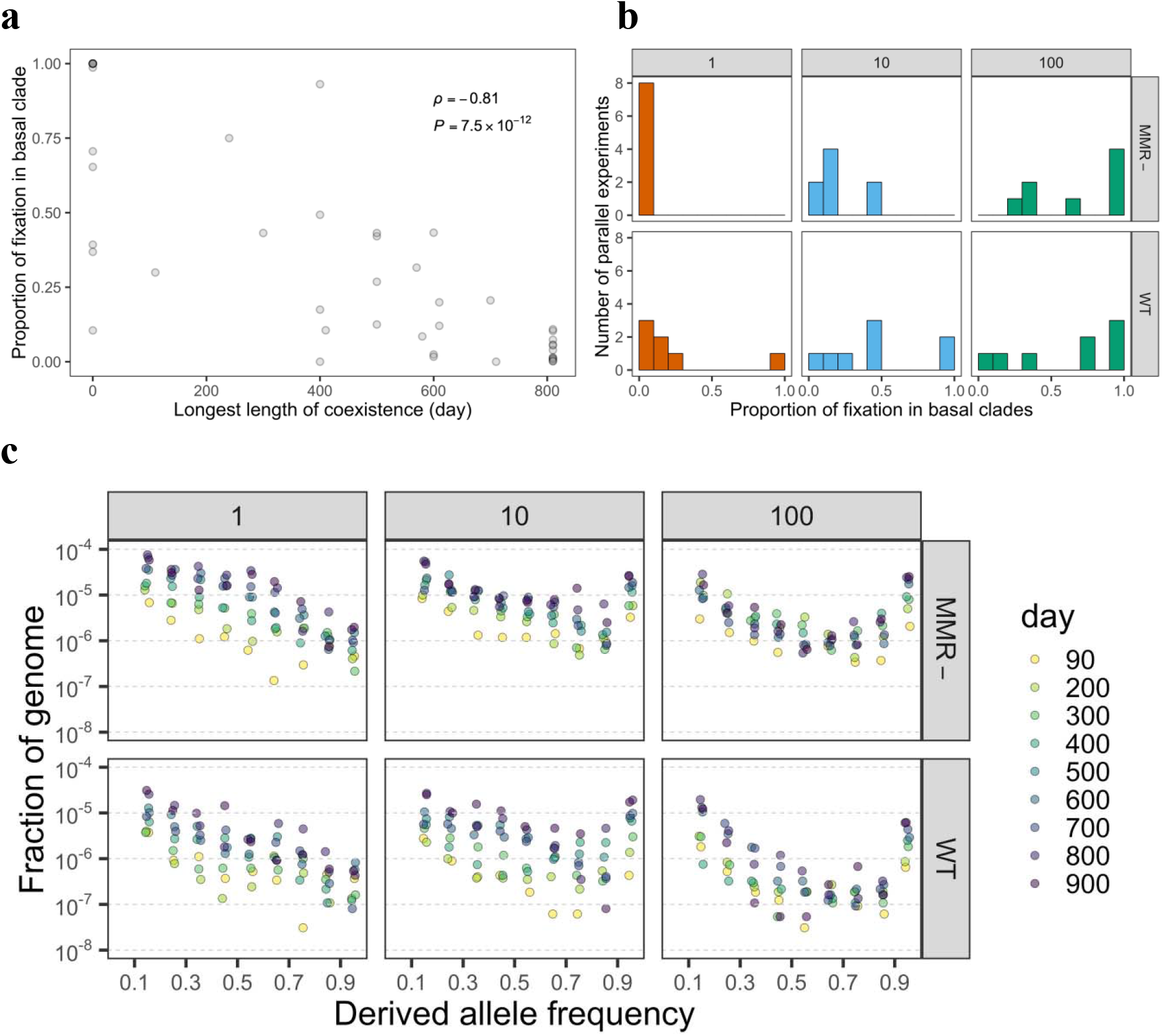
Additional evidence for less population substructure in 100-day populations. **(a)** Negative correlation between the proportion of fixation occurring in basal clades and the longest length of coexistence across all experimental populations. Each dot is an experimental population. *ρ* : Spearman’s rank correlation coefficient. *P*: *p*-value of Spearman’s rank correlation. **(b)** The distribution of proportion of fixation occurred in basal clades for each resource-limitation cycle/genetic background combination. We found resource-limitation cycle length significantly affects the proportion of fixation occurred in basal clades (*P* = 2.5 x 10^−4^, Scheirer-Ray-Hare test). The difference is significant between 1-day and 100-day populations (*P* = 5.5 x 10^−5^, Mann-Whitney *U* test), 10-day and 100-day populations (*P* = 0.016, Mann-Whitney *U* test), and daily and 10-day populations (*P* = 1.6 x 10^−3^, Mann-Whitney *U* test). **(c)** Temporal distribution of mean derive allele frequencies (DAF) in evolved populations. Each dot shows a mean fraction of genome showing a range of DAF with bin size 0.1 in different time point of experimental evolution. The different time points of experimental evolution are shown by colors.

**Supplemental Figure 3.**
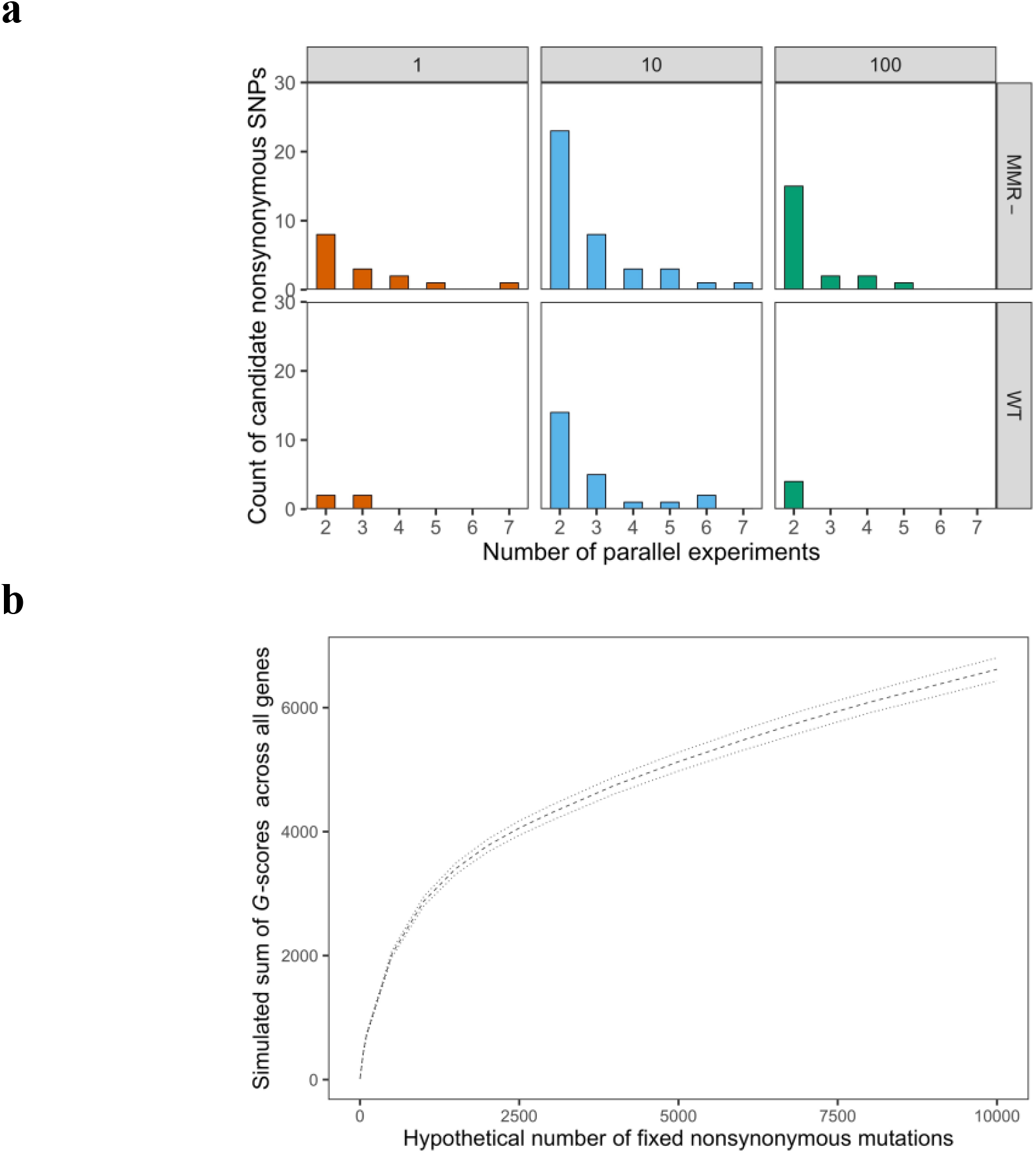
Observed and expected parallelism in experimental evolution. **(a)** shows the nucleotide-level parallelism observed in our experimental evolution. For each combination of a resource-limitation cycle (1 day, 10 days, or 100days) and a genetic background (MMR-for mismatch repair defected, or WT for wild-type), we plotted the distribution of number of parallel experiments where the same fixed nonsynonymous SNPs are identified. Due to a very low probability that any one nucleotide is mutated at least twice (< 10^−4^ by simulation with a random sample of 10^4^ mutations), our observed nucleotide-level parallelism again suggests positive selection in action. **(b)** illustrates that the null expectation of *G*-scores across genes vary with total number of mutations considered. The dashed line and dotted lines summarize the mean and 95% range across 1000 simulations of hypothetical number of fixed nonsynonymous mutations = 1, 50, 100, 500, 1000, 1500, 2000, 2500, 3000, 4000, 5000, 6000, 7000, 8000, 9000, 10000.

**Supplemental Figure 4.**
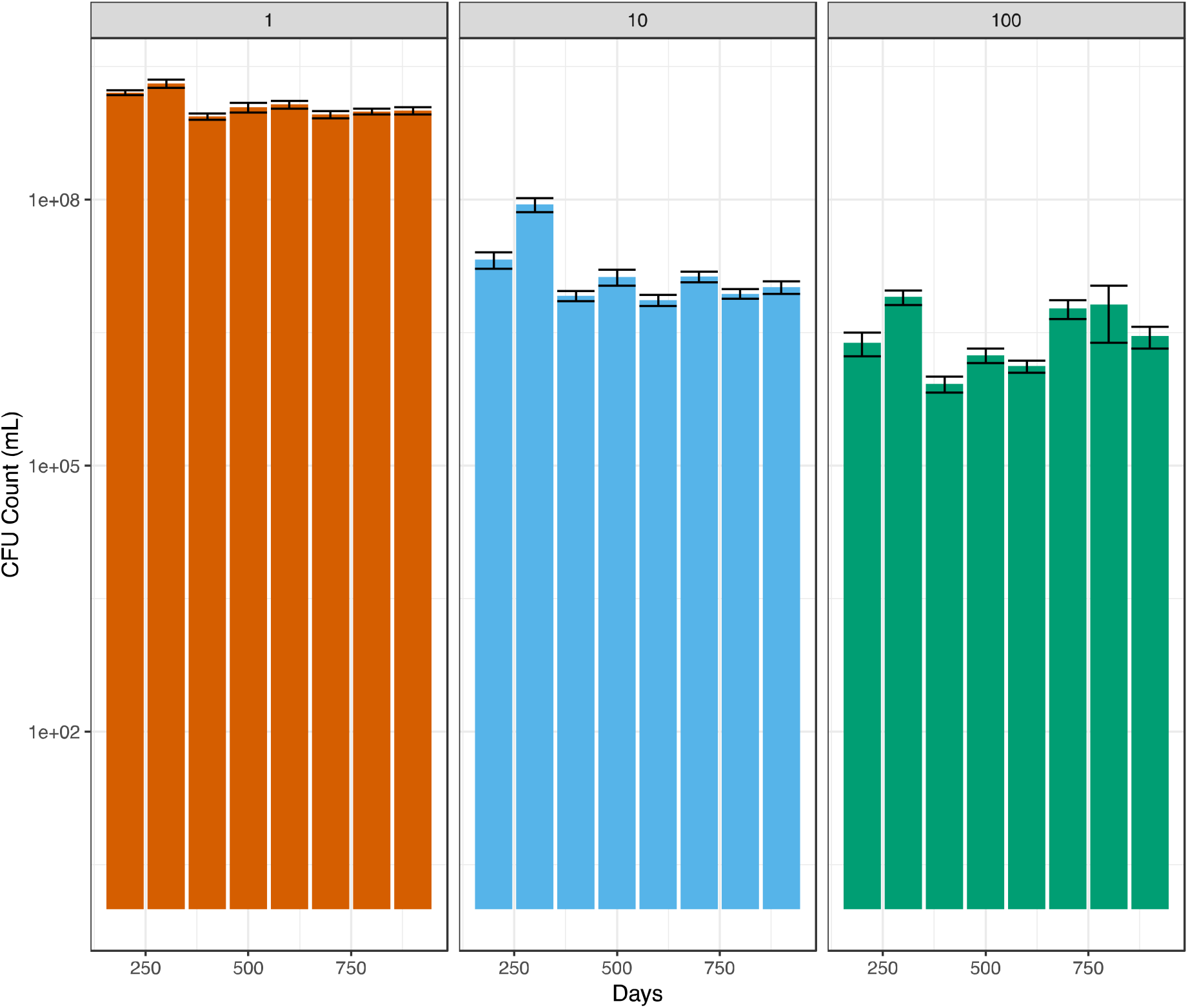
Density measurement of experimental populations during experimental evolution. Pre-transfer population density of 1- (orange), 10- (blue) and 100-day (green) populations as determined by CFU counts. Bars represent mean population density of WT and MMR-populations after 200, 300, 400, 500, 600, 700, 800, and 900 days of evolution and error bars represent ±s.e.m.

**Supplemental Table 1.** List of sequencing profiles used in the analysis

**Supplemental Table 2.** Molecular functions of genes overrepresented for fixed nonsynonymous SNPs in each combination of resource-limitation cycle length and genetic background.

**Supplemental Table 3.** Molecular functions of genes overrepresented for fixed structural mutations in each resource-limitation cycle length and genetic background.

**Supplemental Table 4.** List of candidate mutations found in genes which may alter mutation rates

**Supplemental Table 5.** List of significantly enriched Gene Ontology terms and KEGG pathways of genes with significant G-scores in MMR-populations with 10-day cycles but not in populations with daily cycles.

**Supplemental Table 6.** List of significantly enriched Gene Ontology terms and KEGG pathways of genes with significant G-scores in MMR-populations with 100-day cycles but not in populations with daily cycles.

**Supplemental Table 7.** List of significantly enriched Gene Ontology terms and KEGG pathways of genes with significant G-scores in WT populations with 10-day cycles but not in populations with daily cycles.

**Supplemental Table 8.** List of significantly enriched Gene Ontology terms and KEGG pathways of genes with significant G-scores in WT populations with 100-day cycles but not in populations with daily cycles.

**Supplemental Table 9.** List of fixed nonsynonymous SNPs which hit identical nucleotides in multiple experimental populations for each combination of resource-limitation cycle and genetic background.

